# Measuring the 3D wake of swimming dice snakes using volumetric velocimetry

**DOI:** 10.1101/2022.11.04.515204

**Authors:** Vincent Stin, Ramiro Godoy-Diana, Xavier Bonnet, Anthony Herrel

## Abstract

Experimental observations of the 3-dimensional wake of swimmers are scarce. This study provides the first experimental measurements of the three-dimensional vortical structures of an anguilliform swimmer. A volumetric velocimetry (DDPTV) setup was used to quantify the wake of freely swimming dice snakes (*Natrix tessellata*) in a tank. Multiple swimming trials were recorded with three snakes swimming at a forward swimming speed ranging from 0.2 to 0.8*L*.*s*^−1^. The volumetric measurements show that during swimming, *Natrix tessellata* sheds vortex tubes from different parts of its body with oscillating lateral undulations. The vortex tubes are then linked to form a hairpin-like structure as predicted by computational fluid dynamic analyses. Quantitative measurements show that the vortex size gradually increases over time. The vortex circulation decreases after attaining a maximum. These results provide an experimental demonstration that the 3D wake structure of an anguilliform swimmer takes well organized dynamic shapes as predicted and provide baseline data for future studies comparing the wake structure of snakes with different locomotor ecologies.

## INTRODUCTION

Undulatory swimming kinematics are usually classified in four main modes involving different proportions of the body and/or caudal Fin (BCF) (Lindsey 1978; Sfakiotakis, Lane, and Davies 1999). Anguilliform swimming is traditionally used to describe the swimming of elongated swimmers. The 2D kinematics of anguilliform swimming can be described by an undulation that increases in amplitude along the body. The hydrodynamics of anguilliform swimming have long been studied. Since Lighthill’s analytical Large Amplitude Elongated Body Theory (Lighthill 1971), numerical investigations have been conducted (Battista 2020a,b; Borazjani and Sotiropoulos 2010; Borazjani and Sotiropoulos 2009; Kern and Koumoutsakos 2006; Khalid et al. 2020, 2021; Nangia et al. 2017; Ogunka et al. 2020) in order to obtain a direct estimation of quantities such as forces, pressure fields and swimming efficiency for an idealized swimmer. Anguilliform swimming hydrodynamics have been experimentally studied on eels *Anguilla anguilla* (Muller et al. 2001) and *Anguilla rostrata* (Tytell 2004a,b), lampreys *Petromyzon marinus* (Du Clos et al. 2019; Gemmell et al. 2016; Lehn 2019) and catfish *Plotosus lineatus* (Tack, Du Clos, and Gemmell 2021) using Particle Image Velocimetry (PIV) in 2D.

Although 2D PIV usually offers a good spatial resolution, swimming hydrodynamics needs to be described in three-dimensional space in order to be fully understood as out-of-plane movements are common. Therefore, the use of 3-dimensional particle tracking velocimetry has been used in experimental biology. The main methods are synthetic aperture particle image velocimetry (SAPIV) (Mendelson and Techet 2020), tomographic PIV (Skipper, Murphy, and Webster 2019) and digital defocusing particle tracking velocimetry (DDPTV) (Bartol et al. 2016; Flammang et al. 2011). The main interest of these methods is that they allow to instantaneously capture the whole unsteady flow in three dimensions. However, the three-dimensional wake produced by an anguilliform swimmer has never been characterised experimentally.

The elongated and limbless morphology of snakes is associated with more than ten different gaits used to move in different environments (Jayne 2020). Swimming kinematics were first quantitatively studied by Taylor 1952 inspired by the work of Gray 1933. Subsequent studies (Graham et al. 1987; Hertel 1966; Jayne 1985; Munk 2008) suggested that snakes are anguilliform swimmers. However, previous investigations mostly focused on speed performance (Aubret 2004; Brischoux et al. 2010; Pattishall and Cundall 2008; Wang, Lillywhite, and Tu 2013), or on comparing terrestrial versus aquatic locomotion (Gerald 2017; Isaac and Gregory 2007; Jayne 1988; Shine and Shetty 2001; Shine et al. 2003). Nevertheless, the great diversity of species of different sizes and shapes make snakes suitable candidates to better understand anguilliform locomotion. Moreover, snakes are characterized by several independent radiations into the aquatic environment making them an appropriate group to better understand how elongate limbless tetrapods have adapted their morphology and movements to an aquatic environment.

Here, we provide a first analysis and characterization of the 3D wake structure of an swimming snake. Using DDPTV we measured the wake around free swimming dice snakes (*Natrix tessellata*), a largely semi-aquatic natricine snake that is an excellent swimmer and that feeds nearly exclusively on fish. We document the temporal and spatial changes of the vortical structures by quantifying their morphology, size, and strength. We then compared results to predictions of the wake structure of anguilliform swimmers based on previous computational fluid dynamics and 2D experimental PIV studies.

## MATERIAL AND METHODS

### Animals

Swimming kinematics and PIV data were obtained from three adults captives dice snakes (*Natrix tessellata*). Dice snakes are semi-aquatic snakes and were chosen for their relatively small size.

### Swimming kinematics

In order to characterize their swimming behaviour, 39 swimming trials were recorded (*N*_*M*_ = 17, *N*_*F*1_ = 11, *N*_*F*2_ = 11) with a high speed camera (Phantom MIRO, 1000fps) by filming snakes swimming in a 4m-long, 40cm wide plexiglass swimming tank. Animals were filmed in ventral view and illuminated using two infrared lights. Kinematic parameters (Fig.1) were extracted by digitizing the snake’s midline using an custom-written Matlab routine.

**Fig. 1.**
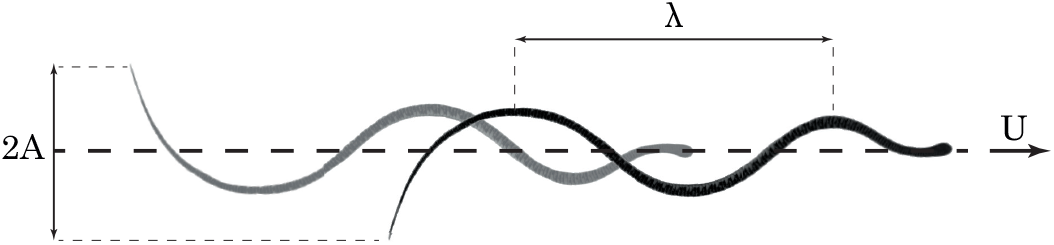
Swimming kinematics. Superimposed images of a snake at two consecutive maxima of tail deflection. The tailbeat amplitude *A*, tailbeat frequency *f* and the forward velocity *U* are measured by finding the peaks in the lateral excursion of the tail-tip over time for at least one full tailbeat cycle. The body wave length *λ* is measured when two adjacent crests are clearly visible on the snake body.

We measured the following anatomical and kinematic parameters: the total length, from the tip of the snout to the tip of the tail (*L*), maximal diameter (*d*), body mass (*w*), forward swimming speed (*U*), tailbeat frequency (*f*), tailbeat amplitude (*A*) and body wavelength (*λ*). These data are summarized in Table 1. The Reynolds and Strouhal numbers were also computed as :

**Table 1.**
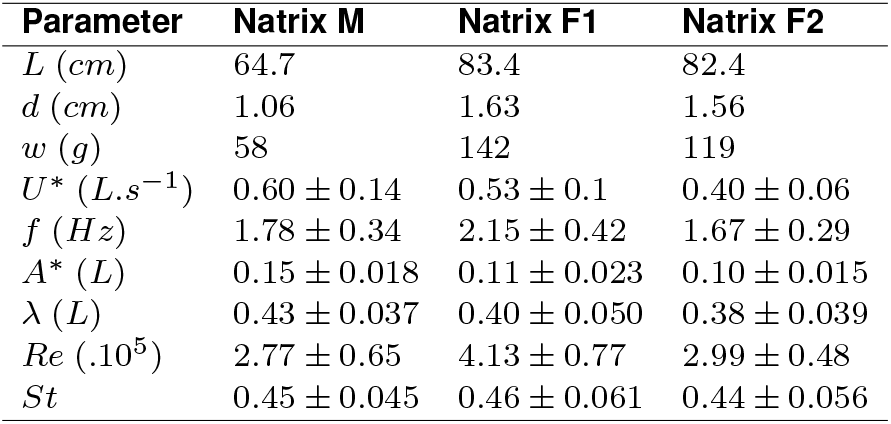
Anatomical & kinematic parameters. Number of swimming trials : *N*_*M*_ = 17, *N*_*F*1_ = 11, *N*_*F*2_ = 11.

| Parameter | Natrix M | Natrix F1 | Natrix F2 |
| --- | --- | --- | --- |
| $L$ (cm) | 64.7 | 83.4 | 82.4 |
| $d$ (cm) | 1.06 | 1.63 | 1.56 |
| $w$ (g) | 58 | 142 | 119 |
| $U^*$ ($L.s^{-1}$) | $0.60 \pm 0.14$ | $0.53 \pm 0.1$ | $0.40 \pm 0.06$ |
| $f$ (Hz) | $1.78 \pm 0.34$ | $2.15 \pm 0.42$ | $1.67 \pm 0.29$ |
| $A^*$ (L) | $0.15 \pm 0.018$ | $0.11 \pm 0.023$ | $0.10 \pm 0.015$ |
| $\lambda$ (L) | $0.43 \pm 0.037$ | $0.40 \pm 0.050$ | $0.38 \pm 0.039$ |
| $Re$ ($\cdot 10^5$) | $2.77 \pm 0.65$ | $4.13 \pm 0.77$ | $2.99 \pm 0.48$ |
| $St$ | $0.45 \pm 0.045$ | $0.46 \pm 0.061$ | $0.44 \pm 0.056$ |

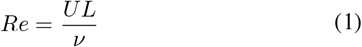

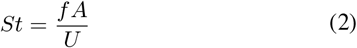

With *ν* the water kinematic viscosity.

Pairwise t-tests show no statistical differences between the 3 snakes for all the measured kinematic parameters (*p*_*U*_ = 0.155, *p*_*f*_ = 0.205, *p*_*A*_ = 0.673, *p*_*λ*_ = 0.282, *p*_*Re*_ = 0.180, *p*_*St*_ = 0.785).

### Experimental PIV setup

In order to quantify the flow around a swimming snake we used a volumetric 3-component DDPTV setup (V3V-9000-CS system, TSI ; Fig.2). The experiments were conducted in a water filled plexiglass tank (30 × 210 × 18 cm). The water was seeded with 50 *µ*m polyamide particles (PSP-50, Dantec Dynamics) with a concentration of around 5.10^−2^*ppp* (Cambonie and Aider 2014). The tank was illuminated from above by a 14 Hz pulsed 200mJ dual head Nd:YAG laser (Quantel Evergreen) expanded with two -25mm cylindrical lenses. The particles were filmed by three CCD arrays (Powerview Plus 4MP-HS) with 50mm camera lenses (Nikon AF Nikkor, aperture f#16) mounted on a 170mm equilateral triangular frame (V3V 9000-CS). The acquisition setup was piloted by the INSIGHT V3V-4G software and calibrated as explained by Troolin and Longmire 2010. The resulting measuring volume is the intersection between the fields of view of the cameras and the laser cone. Its dimension is approximately 18 × 18 × 12cm and it is centered on the water tank in width (Z-axis) and height (X-axis).

**Fig. 2.**
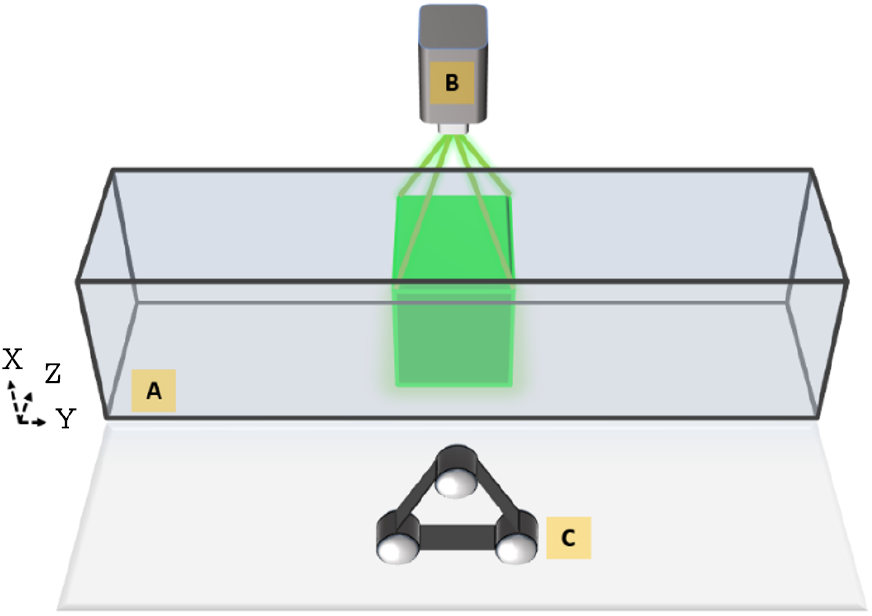
Volumetric imaging setup. A: Water tank, B: Nd:YAG laser (Quantel Evergreen), C : Camera frame

### Recording procedure

During each trial the snake was placed at one side of the tank and allowed to swim. When put into the water snakes typically swam to the other side of the tank along the *X* direction, passing through the measurement volume while being recorded by the cameras at a frequency of 14Hz. Preliminary trials were conducted to estimate the best ΔT between two consecutive PIV frames so that displacement of the seeding particles does not exceed 6 pixels. The resulting ΔT was between 1ms and 2.5ms. The PIV swimming trials were performed on six different days over the course of several weeks. During the first two recording days, females were gravid which impacted their mass and body diameter. The measurements in Table 1 were made after the batches of eggs were laid.

### Data processing

#### Image masking

In order to avoid artefacts induced by the presence of the snake’s body on the raw images during the PIV analysis, the snake was removed from the images with the help of an custom-written masking routine in MATLAB using the Image Processing Toolbox.

#### PIV Processing

The images were then processed using the INSIGHT V3V-4G software in order to obtain the vector fields. The main steps of the data processing are: (1) 2D particle identification in the images of each camera using a Gaussian fitting (Levenberg–Marquardt algorithm). (2) 3D particle identification, the correspondence of a particle in one image to the same particle in the other two images is carried in agreement with the calibration. The fine search tolerance is set to 0.5pxl and the coarse search tolerance to 1pxl. (3) The velocity is processed using the 3D displacement between two consecutive frames with a relaxation algorithm (Pereira et al. 2006). (4) The neighbor tracking reconstruction algorithm uses the trajectories from the neighboring particles to find the probable location of a missing triplet at time *t* + ΔT. (5) The randomly spaced vectors and finally interpolated using a Gaussian-weighted algorithm (Pereira and Gharib 2002).

#### Base change

The vector fields were then imported in an other custom-written MATLAB routine for processing. The swimming trials being in free swimming conditions, the movements of the snake in the measurement volume can be quite different from one sequence to another which makes the comparison challenging. In order to tackle this problem, we changed the regular Cartesian coordinate system (*XY Z*) to a new rotated coordinate system (*X*_*rot*_*Y Z*_*rot*_) for each swimming trial. For each recorded sequence the water jet created by the last tailbeat was tracked with a velocity thresholding method. The trajectory of the jet is estimated by a linear regression of the superimposed position of the jet center. A new base is then created by a rotation around the Y-axis so that the new *X*_*rot*_ axis is co-linear to the jet’s trajectory and the *Z*_*rot*_ axis perpendicular to it.

### Hydrodynamic parameters

The unsteadiness of the flow field induced by the successive tail beats is quantified by computing the Q-criterion of the vector field. Q is defined as the second invariant of the velocity gradient tensor (Hunt, Wray, and Moin 1988):

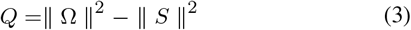

With Ω the antisymmetric part and *S* the symmetric part of the tensor. The positive values of *Q* indicates the parts of the flow field were vorticity dominates over the viscous stress. The strength of the vortices was estimated by computing the circulation of the vortex cores:

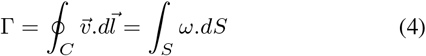

The surfaces *S* of the vortex cores are identified by using the Γ_2_ criterion (Graftieaux, Michard, and Grosjean 2001) in the rotated *X*_*rot*_*Y* planes co-linear to the jet trajectory so that a vortex core is always present in the planes. The vortex size *d* is defined by the distance between the center of the positive and negative vortex cores:

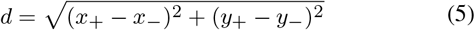

With (*x*+, *y*+) and (*x*−, *y*−) the coordinates of the centers of the positive and negatives vortex cores in the *X*_*rot*_*Y* planes. In order to compare the results between the different individuals and sequences, the hydrodynamic parameters are non-dimensionalized by the mean forward swimming speed *U* of the snake during the measurement period and the snake diameter *D* :

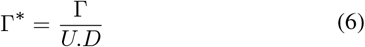

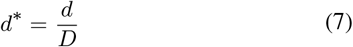

## RESULTS

### Swimming kinematics

A total of 112 sequences of free swimming *Natrix tessellata* with at least one tailbeat in the measuring volume were recorded. The number of sequences were respectively 31, 52 and 29 for Natrix M, Natrix F1 and Natrix F2. One striking feature was the complex three-dimensional displacement of the snake when it swam. The snakes were swimming with the characteristic lateral undulation of an anguilliform swimmers in the frontal plane (XZ) but there were also noticeable albeit smaller oscillations in the sagittal plane (XY). The tail is often positioned below the body and the tail-beats were downward facing. The median mean swimming speed in body lengths per second was respectively *U*_*F*1_ = 0.34*L*.*s*^−1^, *U*_*F*2_ = 0.33*L*.*s*^−1^ and *U*_*M*_ = 0.47*L*.*s*^−1^ (Fig.3). A pairwise t-test shows that there is no statistical difference in the dimensional swimming speed of the 3 swimmers (*p* = 0.28) but there is a high statistical difference (*p* < 10^−8^) in the relative swimming speed of the male compared to the two females (Fig.3). The swimming speed of the two females was significantly lower than in the kinematic experiment but this was not the case for the male.

**Fig. 3.**
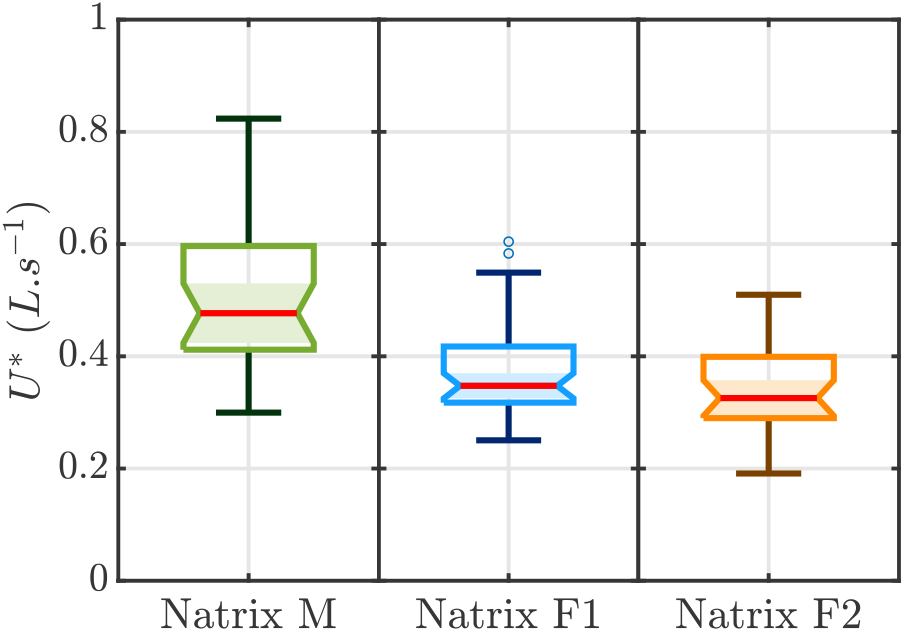
Boxplot of mean voluntary forward velocity of the swimming snakes. (*N*_*M*_ = 31, *N*_*F*1_ = 52, *N*_*F*2_ = 29). The relative forward swimming speed of the smaller male was significantly greater than that of the two females.

### Vortex dynamics

Our analysis shows that vortices were shed after every change of direction of the swimmer’s body during oscillatory movements. The displacement of the body creates two vortex tubes of opposite rotation (Fig.4.A,B) on top and on the bottom which eventually connect to each other. The shed vortices were tilted at an angle of approximately 30 to 40° to the swimming direction (Fig.4.C). Their trajectory was mostly linear with a tendency to curl.

**Fig. 4.**
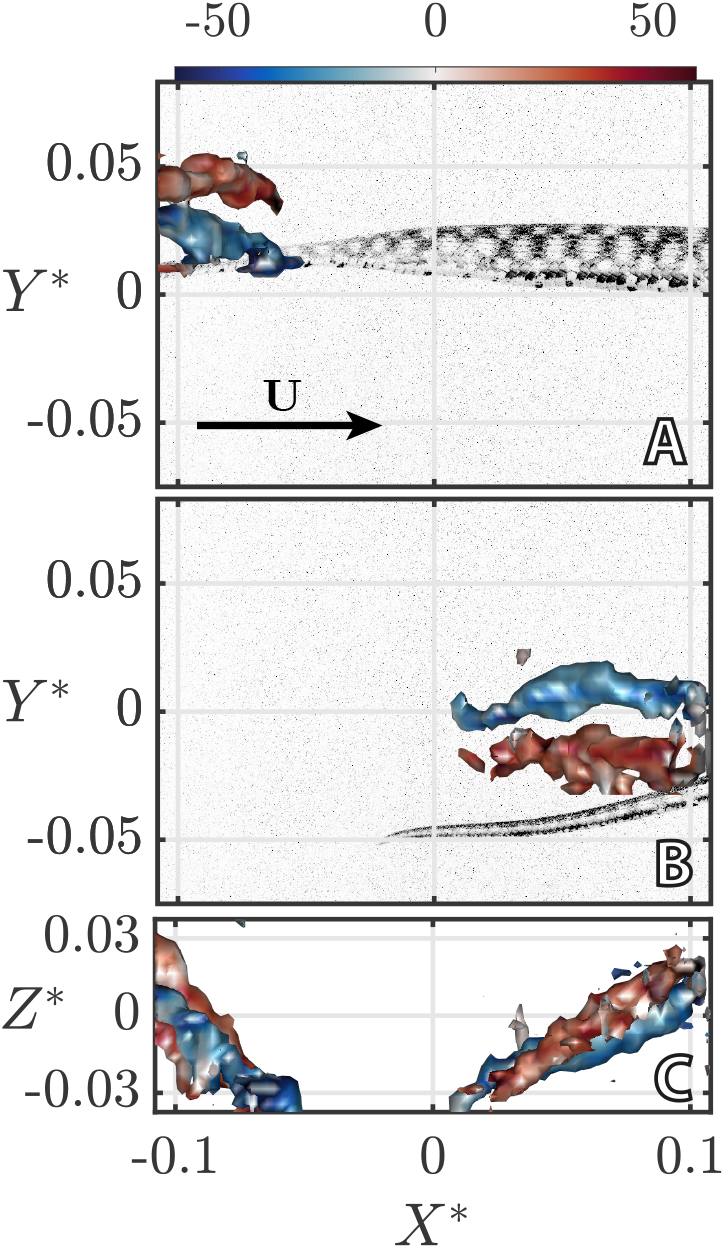
Q-criterion (Q=80) *ω*_*X*_ -colored vortex isosurfaces. (A,B) Side view of instantaneous trailing edge vortex isosurfaces with corresponding snake image (NatrixF1) at two successive time steps (Δ*T* = 286*ms*). (C) Top view of superimposed isosurfaces at their respective times. X,Y and Z axis are non-dimensionalised by the length of the snake. The corresponding swimming trial video with the computed isosurfaces and measured hydrodynamic parameters is available in supplementary.

The spatial and temporal resolution of the setup precluded capturing the birth of the vortex. The vortices were monitored from their detection in the volume until they either reached the edges of the working volume or dissipated. The circulation of the vortex cores rapidly increased in norm just after the tail beat and reaches a maximum before decreasing (Fig.5.A). The positive and the negative vortex cores displayed symmetrical variations during the observed time period. During its lifetime, the vortex size gradually increased (Fig.5.B) and changed from a flattened into a more circular shape.

**Fig. 5.**
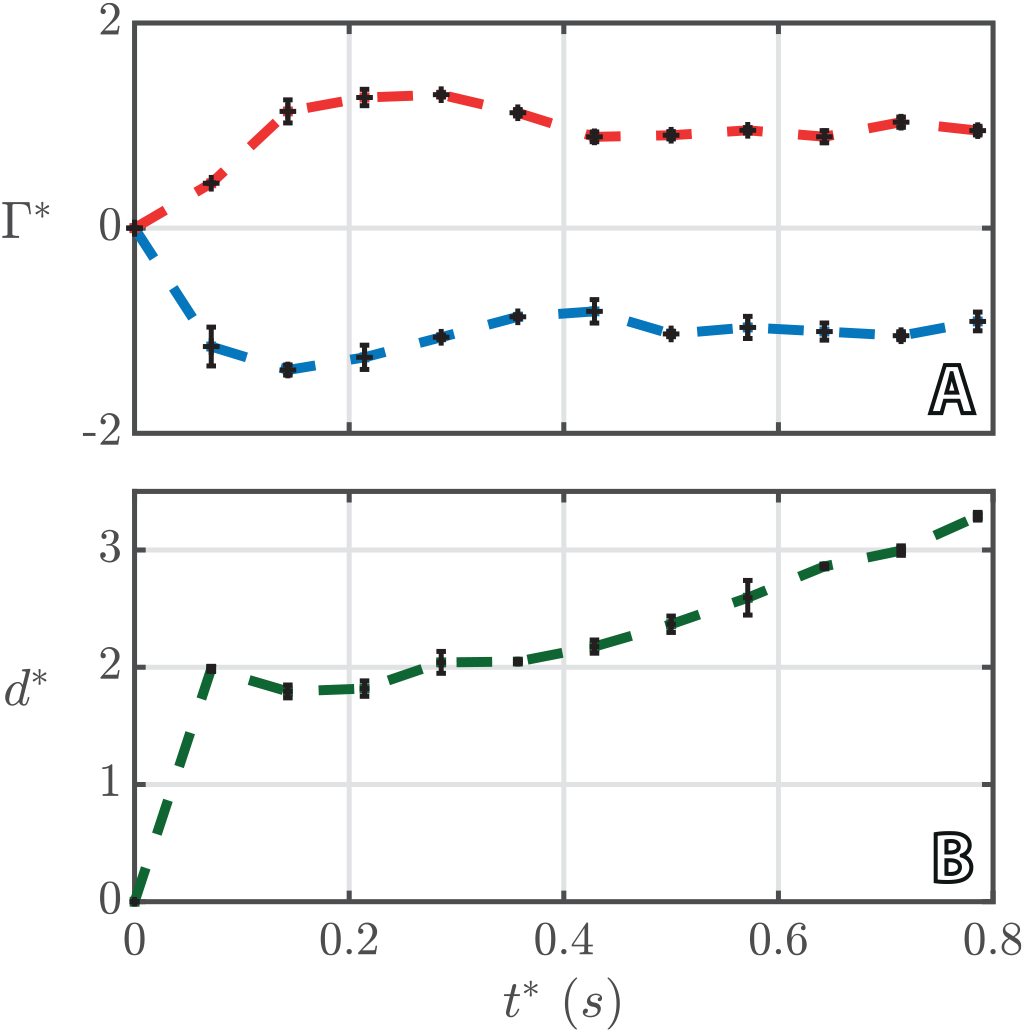
(A) Positive (red) and negative (blue) vortex cores non-dimensionalized circulation of the second trailing edge vortex from the frame before its detection (*t*\* = 0) to the time when it reaches the edge of the measurement volume. (B) Size-adjusted vortex size evolution. Values are the mean values in 3 consecutive planes centered on the vortex path. Error bars are the standard deviations of the values.

### Multiple sequence comparisons

To quantify differences in the hydrodynamic parameters across individuals sequences we focused on trials where at least one tail beat vortex was clearly visible and where measurements were possible (*N*_*M*_ = 12; *N*_*F*1_ = 13; *N*_*F*2_ = 14). Snakes were freely swimming in the water tank and adopted various behaviours. As a result, there was a diversity in the circulation range of subsequent sequences of the same individual. The dimensional maximal circulation ranged from around 18 to 51*cm*^2^.*s*^−1^, 25 to 70*cm*^2^.*s*^−1^ and 22 to 88*cm*^2^.*s*^−1^ for NatrixM, NatrixF1, and NatrixF2, respectively. Most of the non-dimensional circulation curves were contained with a range between 0.5 and 1.5 (Fig.6). A pairwise t-test suggests that the maximum dimensional circulation of the measured vortices was significantly lower for the male (*p* = 0.002) but that there was no significant difference in the non-dimensionalized maximum circulations 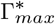 between the three *Natrix tessellate* (*p* = 0.099). The circulation of the vortices produced by the gravid females were the among the highest, especially for NatrixF2. If these swimming trials are removed, sequences from different individuals resemble each other even more (*p* = 0.796). Overall, the circulation either decreases or remains relatively constant over time.

**Fig. 6.**
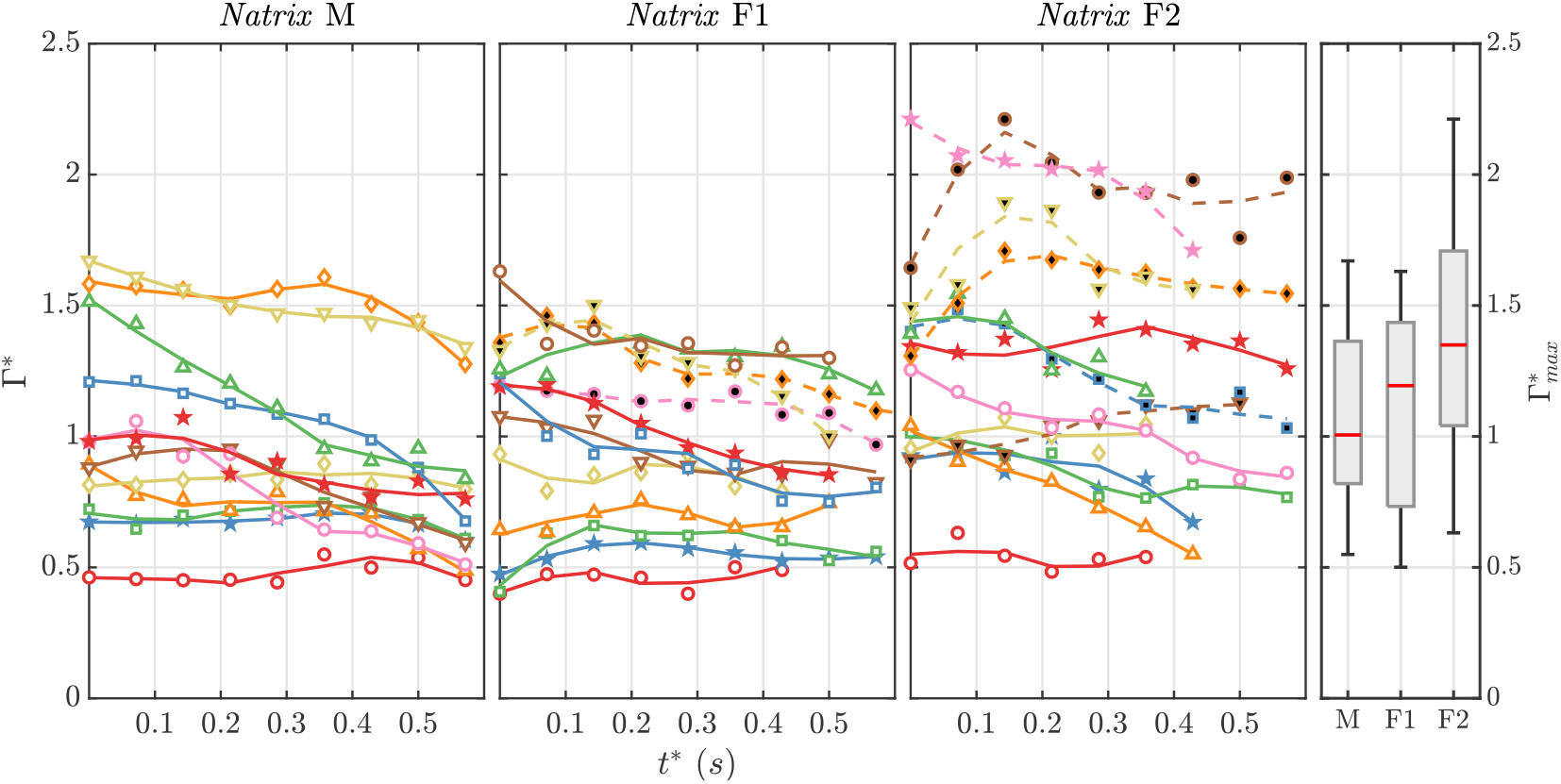
Vortex core circulation statistics for different swimming sequences. *t*\* = 0 correspond to the first frame when the vortex is measurable, meaning that the superimposed plot are not necessarily at the exact same moment of the lifetime of the vortex. Markers are measured data and the line is the moving average smoothing. The dashed plots are the circulations of the vortices produced by gravid females.

The vortex size also varied between sequences. The maximum dimensional vortex size ranged from approximately 2.5 to 5.3*cm*, 2.7 to 5.2*cm* and 2.6 to 5.4*cm* for NatrixM, Natrix F1, and NatrixF2 respectively. There was no significant difference in the maximal vortex size between the three snakes (*p* = 0.824). Similar to what was observed for the vortex circulation, the sizes of the vortices produced by the gravid snakes were the among the largest. The size adjusted vortex size was however, greatest for the male as it has the smallest diameter. (Fig.7). The pairwise t-test (*p* < 10^−4^) suggests that there was a significant difference of the size-adjusted maximum vortex size 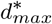 of the male compared to the two females. Overall, the size of the vortices gradually increased or remained relatively constant over time.

**Fig. 7.**
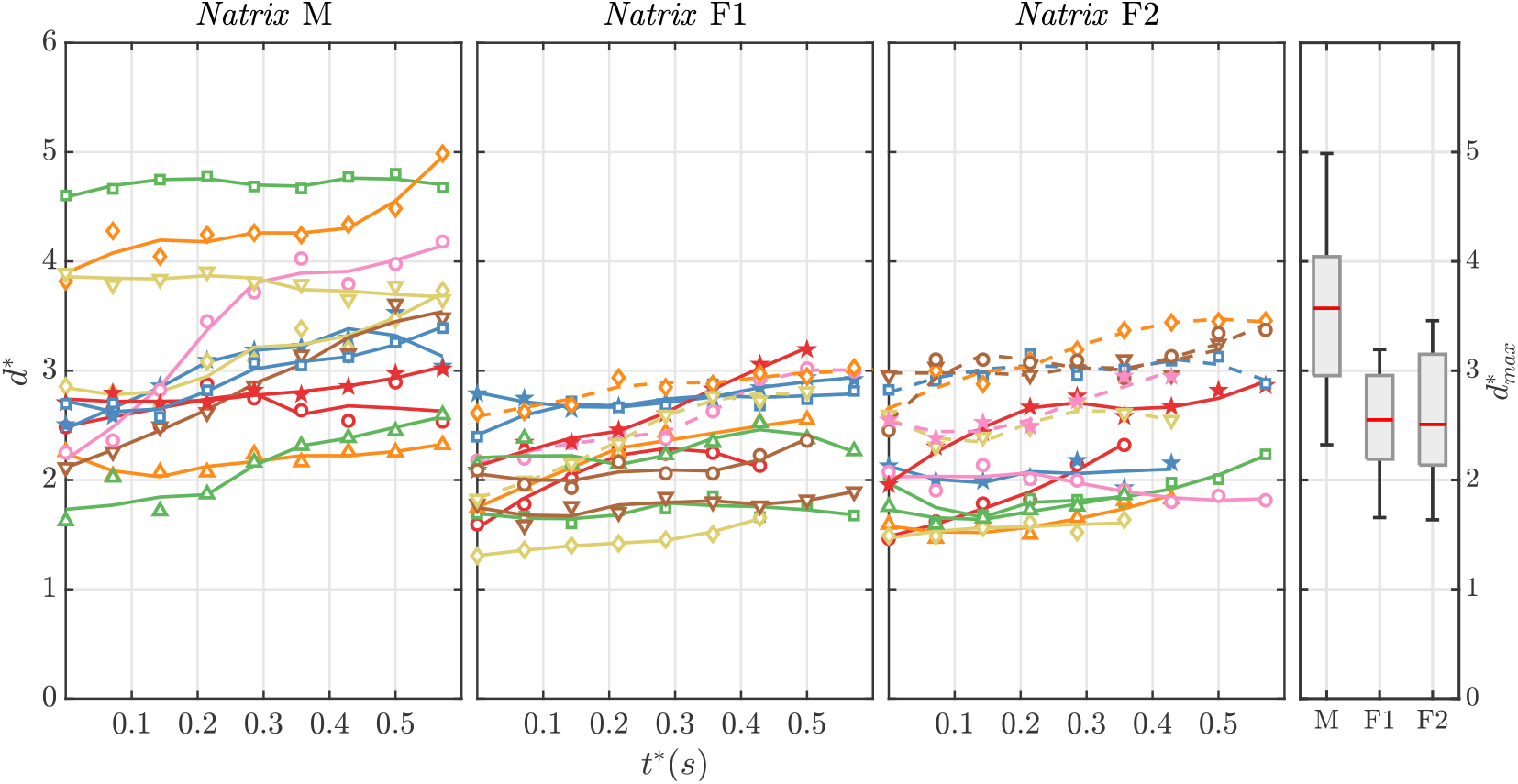
Vortex core size statistics for different swimming sequences. *t* *= 0 correspond to the first frame where the vortex is measurable. Consequently, the superimposed plots do not necessarily represent the same moment in the lifetime of the vortex. Markers represent the measured data and the line is the moving average smoothing. The dashed plots are the vortices produced by gravid females.

## DISCUSSION

### Kinematics

Water snakes are believed to be efficient swimmers by minimizing energy expenditure by the optimization of undulation frequency and amplitude but existing data on snake swimming kinematics are scarce. The Reynolds numbers we obtained were approximately 10 times larger than the ones observed in other anguilliform swimmers (Du Clos et al. 2019; Gemmell et al. 2016; Muller et al. 2001; Tytell 2004b), mostly because of the larger size of the snakes we studied. The mean wavelength was lower than the ones reported for eels (Tytell 2004b) and lampreys (Du Clos et al. 2019). As pointed out by Khalid et al. 2021, a greater wavelength increases thrust but also the frictional drag along the body. The best trade-off between these two quantities depends on the morphology of the swimmer and may result in different kinematic parameters in different animals.

### Vortex morphology

The 2D PIV experiments on eels of Muller et al. 2001 in the frontal plane have led to the prediction that the 3D vortical structure of an anguilliform swimmer should resemble a vortex ring. Tytell 2004b were skeptical about the true roundness of the vortex and the computational work of Borazjani and Sotiropoulos 2009 suggested that the wakes were in fact vortex loops stretched in the stream-wise direction or “hairpin-like” vortices. The traditional plane to measure the swimmer’s wake in the existing literature is the frontal plane (XZ) without looking at the other ones (Fig.8.A,B,C). Our experimental results in the frontal plane show similar results with two vortex cores on the sides of a lateral jet but the 3D morphology conformed to predictions generated by computational studies with connected vortex tubes alternating their signs every tail beat. Our results show that whereas the vortex structure is stretched in the longitudinal direction, a major part of the rotation of the vortex happens in the transverse plane (YZ). A three-dimensional description is therefore needed to fully capture the behaviour and dynamics of the vortices.

**Fig. 8.**
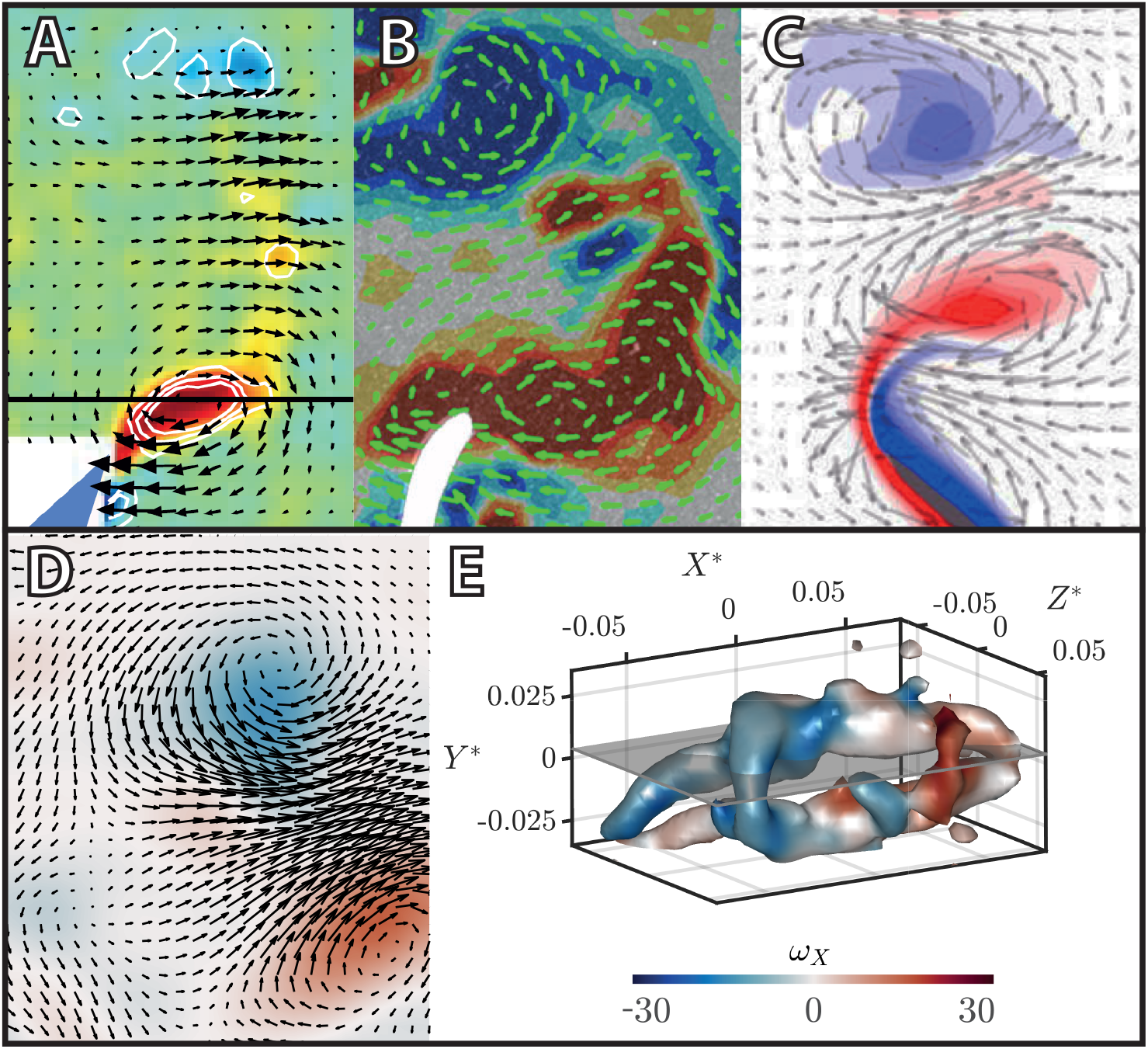
2D vortex comparison between studies. (A,B,C,D) Instantaneous frontal plane (XZ) vector fields of the vortex cores shed by a tail beat of a swimmer for (A) an eel (Tytell 2004b) (B) a lamprey (Gemmell et al. 2016) (C) a CFD model (Kern and Koumoutsakos 2006) and (D) our results. The flow field is smoothed with a 3D Gaussian filter (*σ* = 1.25). The corresponding *ω*_*X*_ -colored 3D plot (E) suggests that the vortex structure is more complex than it appears in a 2D plane. The grey plane is the location of the velocity field (D).

This study focused on the formation mechanisms of the hairpin-like vortices (Fig.9). Just after a tail beat, two vortex tubes of opposite rotation are shed in the wake. The distance between the two vortex tubes scales with the snake’s diameter. As the tail has an elongated cone-like shape, the distance between the two vortex tubes as well as the local circulation gradually increase from the tail tip to the body. The interaction between the two vortex tubes creates a bridge in the region where the circulation / vortex tubes distance ratio is the highest. The bridge creates a connection of the two vortex tubes and a detachment of the vortex tubes before the bridge. From the vortex dynamics point of view, theses steps are similar to the events described by Zhou et al. 1999 for coherent packets of hairpin vortices in a channel flow. The rounding of the vortex structure after the bridging step is also reminiscent of the Ω-shaped vortex created after the self induction of the hairpin vortex. The bridging step is fast so with our relatively slow framerate it is easy to miss but almost impossible to fully describe using 2D settings.

**Fig. 9.**
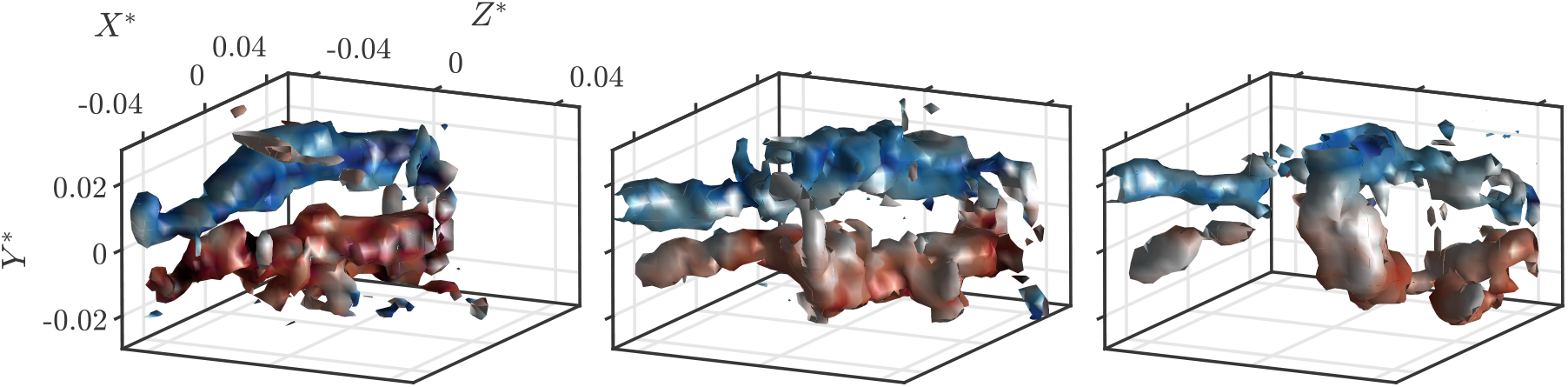
Main steps of the behaviour of a shed vortex. *ω*_*X*_ -colored isosurfaces of Q-criterion (*Q* = 60) (A) Two vortex tubes of opposite rotation (*t*\* = 71*ms*). (B) Bridging of the two vortex tubes (*t*\* = 285*ms*). (C) Separation of the tubes before the bridge and the shape of the structure is getting rounder (*t*\* = 500*ms*). The mean vorticity at the bridge section (*ω*_*Y*_ = 15.08*s*^−1^) is similar to the vorticity of the vortex tubes (*ω*_*X*_ = 15.45*s*^−1^).

Vortex behaviour was also witnessed more anteriorly for the body vortices. When the body wave attains a maximum and then changes its direction a body vortex is created as predicted by simulations (Khalid et al. 2021; Li et al. 2016) and observed in 2D experimental studies of anguilliform swimmers (Gemmell et al. 2016; Muller et al. 2001). Just after the change of direction, two parallel vortex tubes are shed in the wake and then connect, as observed for the vortices produced at the tail tip. Our cameras film from the side of the snake so body vortices were often partly masked as they were most of the time aligned with the body of the snake. The production of body vortices has been described by Gemmell et al. 2016 on lampreys where a particular attention was given to the bending of the body. On top of the oscillations, the body has an angular velocity which helps to rotate the fluid along the body and creates adjacent vortices. Although this was not measured in our study, the bending of the snake’s body was observed in our video recordings suggesting that the angular velocity of the snake’s body plays a role in the creation of body vortices in addition to the change of direction of its oscillation. The production of counter-rotating vortex tubes along the body has been suggested to generate additional trust (Fu and Liu 2015) during swimming.

### Vortex size and circulation

The hairpin-like vortices were characterized by their circulation and size. The absolute maximal circulation measured for the three swimmers was similar to the ones measured in smaller anguilliform swimmers : ∼ 20*cm* eels (Tytell 2004b) and ∼ 10*cm* lampreys (Gemmell et al. 2016). The circulation of small hairpin vortices decreases exponentially after attaining a maximum because of the viscous interactions of the counter rotating vortex tubes (Sabatino and Maharjan 2015). On the other hand, the circulation of laminar vortex rings slowly decreases after the maximum (Rosenfeld, Rambod, and Gharib 1998). In our results, depending on the swimming trial, the circulation either decreases or stays relatively constant. The decreasing vortex circulation is probably due to the viscous interactions with either the counter rotating vortex tubes or surrounding vortices created by the other tail beats. The relatively constant or slowly decreasing circulation, on the other hand, behaves like a laminar vortex ring.

Our study also reports data on the evolution of the distance between the two counter rotating vortex tubes created by the oscillations of the anguilliform swimmer. These data are new considering that other anguilliform studies were set in the frontal plane and therefore had no information on the height of the vortex. Studies on vortex rings show that the diameter of the vortex rapidly increases, attains a maximum and then asymptotically decreases to a constant value (Arakeri et al. 2004). The size of the vortices follows the same pattern as they either increase or stay relatively constant. This means that the vortices with an increasing size are in the first part of their lifetime and the ones with a relatively constant size must be near the end of their lifetime.

The two females snakes were recorded before and after they laid eggs. Gravid snakes had a larger body diameter and higher mass compared to non-gravid ones; but proportionaly less locomotor muscles contribute to their overall volume (the eggs represent a load). The load represented by the eggs may impact locomotor perfomance (Seigel, Huggins, and Ford 1987). One of the associated side effects is that gravid snakes typically fast (Lourdais, Bonnet, and Doughty 2002). In our experiments, the gravid snakes shed vortices with a circulation and size that were among the highest recorded, suggesting that they were forced to produce an elevated swimming effort. This may indicate a higher cost of travel and as such gravid snakes likely avoid to forage.

## CONCLUSION

This study provides the first three-dimensional experimental data on the vortical structure produced by an anguilliform swimmer.

With the DDPTV setup we were able to show that free swimming snakes create vortex tubes of opposite rotation at different parts of its body whenever the body wave changes its direction. The vortex tubes then link to form an hairpin-like vortex structure. Theses observations are in agreement with the predictions from computational fluid dynamics studies of anguilliform swimmers. Our measurements made it possible to observe the evolution of the vortex circulation and size over time which are in accordance with existing work on vortex dynamic. Our results lay the foundation for further comparisons such as the interspecific variations of the vortices depending on the morphology and the lifestyle of the swimmer.

## Acknowledgements

This research was supported by Sorbonne University’s Interdisciplinary Doctoral Program (IPV) and ANR DRAGON2 (ANR-20-CE02-0010).

## Contribution

A.H., X.B. and R.G.D. conceived the study; V.S. carried out experimental work; V.S. carried out data analysis; V.S. wrote the manuscript; A.H., X.B. and R.G.D. revised the manuscript. All authors gave final approval for publication.

## Data availability

All data and codes are available from the authors upon request.

